# Adult zebrafish anaesthesia: a study of efficacy and behavioural recovery of different anaesthetics

**DOI:** 10.1101/2021.02.23.432432

**Authors:** Sara Jorge, Jorge M Ferreira, I Anna S Olsson, Ana M Valentim

**Affiliations:** i3S - Instituto de Investigação e Inovação em Saúde, Universidade do Porto, Portugal; Laboratory Animal Science, IBMC - Instituto de Biologia Molecular Celular, Universidade do Porto, Porto, Portugal; Centre for the Research and Technology of Agro-Environmental and Biological Sciences (CITAB), University of Trás-os-Montes and Alto Douro (UTAD), Vila Real, Portugal

**Keywords:** anaesthesia, recovery, novel tank, zebrafish

## Abstract

The use of proper anaesthesia in zebrafish research is essential to ensure fish welfare and data reliability. However, anaesthesia long-term side effects remain poorly understood. The purpose of this study was to assess anaesthesia quality and recovery in adult zebrafish using different anaesthetic protocols and to determine possible long-term effects on the fish activity and anxiety-like behaviours after anaesthesia.

Mixed sex adult AB zebrafish were randomly assigned to 5 different groups (control, 175mg/L MS222, 45 mg/L clove oil, 2 mg/L etomidate and 5mg/L propofol combined with 150mg/L lidocaine) and placed in the respective anaesthetic bath. Time to lose the equilibrium, response to touch and to tail pinch stimuli, and recovery after anaesthesia administration were evaluated. In addition, after stopping anaesthesia, respiratory rate, activity and anxiety-like behaviours in the novel tank test were studied.

Overall, all protocols proved to be adequate for zebrafish anaesthesia research as they showed full recovery at 1h, and only etomidate had minor effects on fish behaviour in the novel tank, a validated test for anxiety.

## Introduction

Versatility is a key characteristic of zebrafish that makes it a popular model in biomedical research, but the development of refinement measures is not always accompanying that popularity and increase in zebrafish use. This includes the development of anaesthetic protocols which are essential to mitigate stress and/ or pain when fish undergo surgical and invasive or stressful procedures.

Tricaine methanesulfonate (MS222), benzocaine, 2-phenoxyethanol, clove oil, etomidate and lidocaine have been described to be used to anaesthetize adult zebrafish, with MS222 use highlighted^1^. From these anaesthetics, clove oil and etomidate were described to cause minimal aversion to zebrafish^2, 3^.

Nevertheless, MS222 is the main sedative and anaesthetic used for zebrafish. Its wide use in this species is probably related to its wide use in other fish, as it is the only anaesthetic authorized for some aquatic species by the US Food and Drug Administration (FDA)^4^. MS222 acts mostly by blocking sodium currents in the nerve membranes, reducing action potentials that leads to muscle relaxation^4-6^. It is a local anaesthetic that acts systemically by administration in a water bath.

Etomidate is an ultra-short acting non-barbiturate hypnotic agent^7^ that provides a rapid induction, but a rather long recovery^8^ and it has no analgesic component^9^.

Clove oil is a local anaesthetic, but, as MS222, it acts systemically when administered by immersion. It is a natural essential oil, the active substance of which is eugenol^10^. Clove oil is highly lipophilic and quickly absorbed through the gills and skin^11^, thus it has rapid induction and delivers consistent anaesthesia in fish compared to other anaesthetics^12^.

As these agents are delivered in water bath, there is an increased risk of overdose and subsequent death or other side effects is high. The use of anaesthetic combinations potentiates anaesthesia, decreasing the risk of overdose by using lower concentrations. One example is the combination of propofol with lidocaine tested by our group which showed promising results^12, 13^, while each anaesthetic alone presents some concerns. Lidocaine hydrochloride, a water-soluble local anaesthetic agent^8^, provides rapid anaesthesia induction and recovery^9^, but exhibits low margin of safety in zebrafish^14^, while propofol, a short-acting sedative-hypnotic used for induction and maintenance of general anaesthesia^8^, does not provide satisfactory analgesia for painful procedures^13^.

In studies reporting anaesthesia parameters and recovery quality, side-effects are often disregarded, overlooking potential consequences for late mortality and/ or interference with research outcomes and fish welfare.

Thus, this study aims not only to assess the efficacy of zebrafish anaesthesia, but also to study the recovery profile of each anaesthetic agent (MS222, clove oil, etomidate and propofol/lidocaine) and their potential to cause long term influence on zebrafish activity and anxiety levels, using the novel tank test.

## Material and Methods

### Ethics statement

All procedures were carried out under personal and project licenses approved by the National Competent Authority for animal research (Direção-Geral de Alimentação e Veterinária) (approval number: 014703), and by the Animal Welfare and Ethics Review Body of the Institute for Research and Innovation in Health for a larger project where this study protocol was included. All experimental procedures were performed in accordance with the European Directive 2010/63/EU on the protection of animals used for scientific purposes, and its transposition to the Portuguese law, ‘Decreto Lei’ 113/2013.

### Animals and Housing

Seventy-five 13 to 16 months old mixed-sex AB zebrafish bred in the Animal Facility of the institute were used. They were kept in 3.5L tanks in a recirculating water system connected to a central unit of water purification and controlled temperature (27°C± 0.2), pH (7± 0.2) and conductivity (900µS) under a 14:10 h light: dark cycle. Fish were fed twice a day with the commercial diet ZEBRAFEED (Sparos, Olhão, Portugal). Food restriction was applied 24 hours before the experiment. After anaesthesia administration, the animals recovered individually for 24 hours in a 1L tank with water at 27 ± 0.5°C, and in visual contact with the neighbours.

During the experiment, the animals were monitored 1, 2, 6 and 24 hours after anaesthesia or longer if they had not recovered normal behaviour.

### Anaesthesia and recovery

Zebrafish were randomly assigned to 5 different groups: Control (unanaesthetized animals; n= 14; pH= 6.94), MS222 (animals anaesthetized with 175mg/L of tricaine methanesulfonate, Sigma-Aldrich, USA; pH= 7.07; n= 15), CO (animals anaesthetized with 45 mg/L of clove oil, Sigma-Aldrich, USA; pH= 7.15; n= 16), Eto (animals anaesthetized with 2 mg/L of 2% etomidate, Lipuro, B. Braun Melsungen AG, Germany; pH= 6.97; n = 15), or P/L (animals anaesthetized with 5mg/L of 2% propofol combined with 150 mg/L of 1% lidocaine, Braun, Queluz de Baixo, Barcarena, Portugal; pH= 7.05; n= 15).

Except for MS222 and clove oil, anaesthetic solutions were freshly prepared. For MS222, a stock solution was previously prepared by adding tricaine methanesulfonate powder to system water and buffering it with sodium bicarbonate to a pH of 7. A stock solution of clove oil 10% was also previously prepared with ethanol.

Anaesthetic baths were prepared in a 1L tank with 200mL of water from the system and then vigorously stirred^13^. Individually, zebrafish was immediately placed in the prepared water bath and time to lose equilibrium, and the reflex to a tail pinch were measured. Equilibrium loss was considered when fish stayed more than 5 seconds in dorsal recumbency, and the response to a tail pinch was observed by gently pressing the caudal fin with forceps. Stimuli were tested approximately every 30 seconds. When 7.5 minutes had elapsed after the loss of equilibrium, the animal was placed in another 1L tank with clean water, and time to regain equilibrium was measured. This recovery was video recorded to assess the opercular movements/ respiratory rate (RR) after the animal was placed in the recovery tank. Control animals were left in a 1L tank with 200 ml of system water without anaesthetics for ∼1 minute to mimic the time spent by treatment groups until loss of equilibrium, and then placed in a similar recovery tank.

One, 2, 6 and 24 hours after the anaesthesia administration, animals were video recorded during 10 minutes by a side-view camera for later analysis by a researcher blinded to the treatments. After recording, a plastic pipette was introduced inside the tank in the field of vision of the fish, and then a light touch with the pipette was applied on the fish side to assess response to visual and mechanical stimuli, respectively. Responses were considered positive when fish moved away from the pipette, increasing activity, freezing or when other behavioural alteration was observed. Also, food was provided at the end of the first period of recording, and fish acceptance was noted. For the video analysis, distance, average speed (m/s), maximum speed (m/s), angular velocity (absolute turn angle (°) / test duration (s)), time (s) and frequency of immobility and freezing were recorded. Also, distance (m) and time (s) spent swimming at a velocity higher than 0.05-0.07 m/s (to roughly assess erratic movements automatically^15^) were evaluated.

To analyse all these parameters, the Any-maze™ behavioural tracking software (Stöelting, Dublin, Ireland) was used. In the software analysis, immobility was considered when the animal was immobile for more than 2 seconds with a sensibility of 70%, while freezing was detected after 1 second of total absence of movements with a threshold of 20.

### Novel tank test

The novel tank test has been used to assess anxiety-like behaviours in fish. It is expected that animals spent more time in the bottom of the tank than in the upper zone, near water surface, from where dangers may appear^16^. The apparatus consisted in a trapezoidal tank (3.5L, Tecniplast) with clean water from the fish system (column of water of 12 cm). Twenty-nine hours after the anaesthesia administration, fish were individually placed in the middle of the novel tank and allowed to explore for a period of 6 minutes. Fish behaviour was video recorded by a side-view camera and the videos were analysed with Any-maze software. In this software, the water column in the tank was virtually divided by a horizontal line in an upper (UP) and bottom (BTM) zone of equal height. Several parameters were assessed in the whole apparatus: distance (m); average speed (m/s); maximum speed (m/s); angular velocity (° s^-1^); time (s), frequency of immobility and freezing (s); distance, time spent and number of movements with fish swimming at a velocity higher than 0.05-0.07 m/s (indicative of erratic movements); immobility and freezing were detected with the same methods described above. Except for angular velocity and indicators of erratic movements, the parameters were analysed in each zone. The latency to enter and exit BTM, number of entries in UP, and the index of distance and time spent in the BTM were also evaluated. These indices were calculated by dividing the distance or time spent in the BTM by the total distance or time, respectively. An index value near 1 means that the animal swam more (distance) or spent more time in the BTM. The same videos of 6 minutes were analysed by dividing that period in two segments of 3 minutes, evaluating the same parameters. The first three minutes may reveal anxiety-like behaviours that can be diluted in the 6 minutes analysis. Comparing the two segments will allow to observe the evolution of behaviour and habituation to the environment.

### Animal numbers

Sample size calculation was performed in G*Power 3.1 (University of Düsseldorf, Germany), assuming type II error probability of α=0.05, a power of 0.90, and an effect size of 0.47. One animal of MS222 and CO were excluded from the recovery analysis because the videos were corrupted. Also, the software could not always detect the fish in one MS222, two Control, four Eto and three P/L videos, being the animals excluded. Moreover, two animals from the MS222 and CO groups and one animal from the Eto group were excluded at the respiratory rate analysis due to poor video quality to measure these movements. Two MS222 and one CO animals were excluded from the novel tank analysis due to corrupted video files, and one control due to lack of animal detection by the software.

### Statistical analysis

Data was checked for normality (Shapiro–Wilk test) and homogeneity of variance (Levene’s test). Normally distributed data is expressed as mean ± standard deviation and non-normally distributed data as median and interquartile range. To assess differences between groups in the anaesthesia measures, one-way analysis of variance (ANOVA) followed by the adequate post-hoc test (Tukey’s or Games-Howell) or Kruskal-Wallis nonparametric test followed by Dunn’s multiple comparison test was used. To analyse distance travelled during anaesthesia recovery, repeated measures ANOVA was used with treatment as between-subjects factor and time as within-subjects factor. Likelihood ratio was used to test if there was an association between the number of animals to react to visual and touch stimuli and the treatment groups in the recovery period.

For novel tank data, paired Student’s t-test or Wilcoxon signed rank were used to assess differences of parameters between the zones and between time segments for each treatment. The One Sample T-test or One Sample Wilcoxon Signed Rank Test were performed to compare the percentage of time spent in each zone with the chance level percentage of time (50%). The same tests were used to compare the angular velocity of each group with the value of 180° s-1, as values above 180° s-1 indicates the presence of erratic/ escape turns^17^.

Data was analysed using SPSS 24.0 (IBM SPSS Statistics 25 Software, IBM Corp, Armonk, NY) and graphical representations were created in GraphPad Prism 7 for Windows (GraphPad Inc., San Diego, CA, USA).

## Results

### Anaesthesia efficacy and recovery

Fish exposed to clove oil lost equilibrium faster than propofol/lidocaine (*p*=0.002; Fig. 1A) treated animals, while MS222-treated animals regained equilibrium faster than all the other groups (*p*<0.0001; Fig. 1C).

**FIG. 1.**
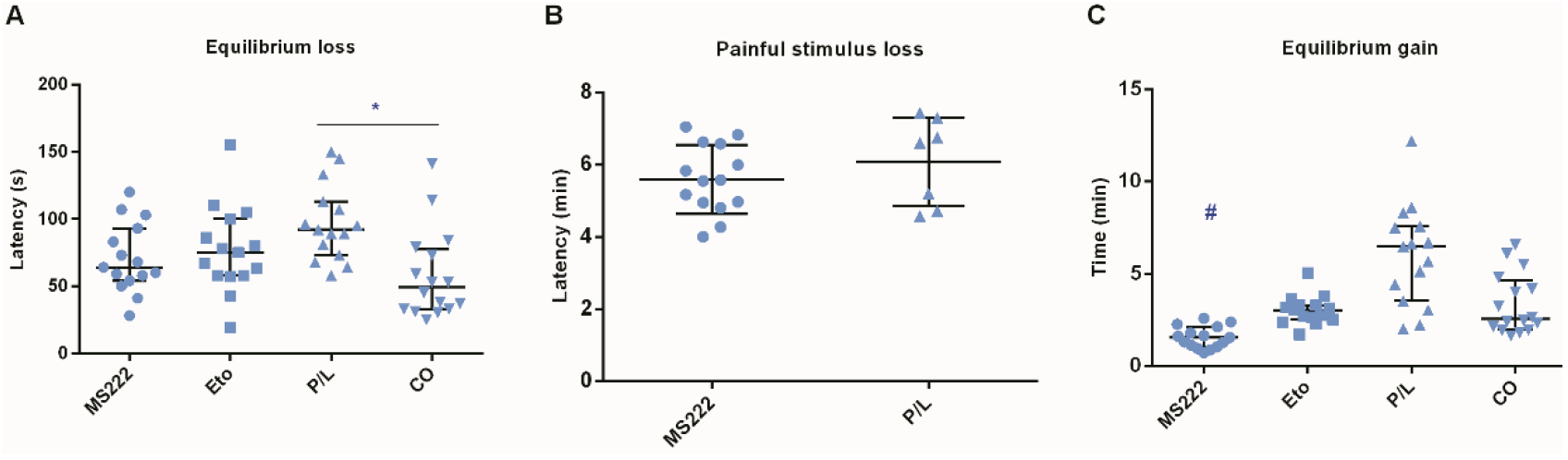
Anaesthetic parameters of adult zebrafish exposed to different anaesthetic protocols (175mg/L of MS222 (MS222); 2 mg/L of etomidate (Eto); 5mg/L of propofol combined with 150 mg/L of lidocaine (P/L); 45 mg/L of clove oil (CO)). **(A)** Time to equilibrium loss. **(B)** Time to loss of reaction to a painful stimulus; **(C)** Time to zebrafish to recover equilibrium after being placed in an anaesthetic bath. Each point represents an animal. In 1A and 1C: n= 15 for all groups, except CO with n= 16; in 1B: MS222 n=14 and P/L n=7. Data are expressed as median [interquartile range]. * p < 0.05; # p≤ 0.05 for comparison between MS222 and all the other groups.

Loss of the pain reflex within the observed time frame varied widely between treatment groups. None or very few animals in the Eto (0/15) and the CO (2/16) groups, about half in the P/L (7/15) and nearly all in the MS222 (14/15) groups lost the pain reflex within 7.5 minutes after equilibrium loss. Nevertheless, there were no differences in the time to lose this reflex (Fig. 1B) when comparing the animals that lost this reflex in P/L (n= 7) and MS222 (n= 14) groups.

All animals responded to the visual and touch stimulus at 2, 6 and 24 hours post-anaesthesia. Immediately after being placed in clean water, only 5 MS222 animals and 3 P/L, CO and Eto animals per group reacted to the visual stimulus. There were no differences between groups regarding the reaction or not to visual or touch stimuli during recovery. Also, there was no association between treatments and number of animals that reacted to each stimulus using the Likelihood ratio.

Regarding swimming distance, repeated measures ANOVA (Supplementary Figure S1) showed a significant time effect (*p*<0.001), with animals swimming longer at 2 hours compared to the 1 hour post-anaesthesia and at 24 hours compared to 6 hours post-anaesthesia. At 1 hour, P/L animals showed higher immobility than the Eto animals (*p*=0.037). At 24 hours, animals exposed to etomidate presented a higher swimming speed than the MS222 animals (*p*=0.045), and the Control animals displayed a decrease in the maximum speed compared with the treatment groups (*p*<0.001).

Concerning the respiratory rate (Fig. 2), P/L (*p*=0.008) and Eto (*p*=0.002) animals had a reduced number of opercular movements compared to MS222 animals. Also, CO animals showed increased respiratory rate compared to those treated with propofol/lidocaine (*p*=0.006) and etomidate (*p*=0.001).

**FIG. 2.**
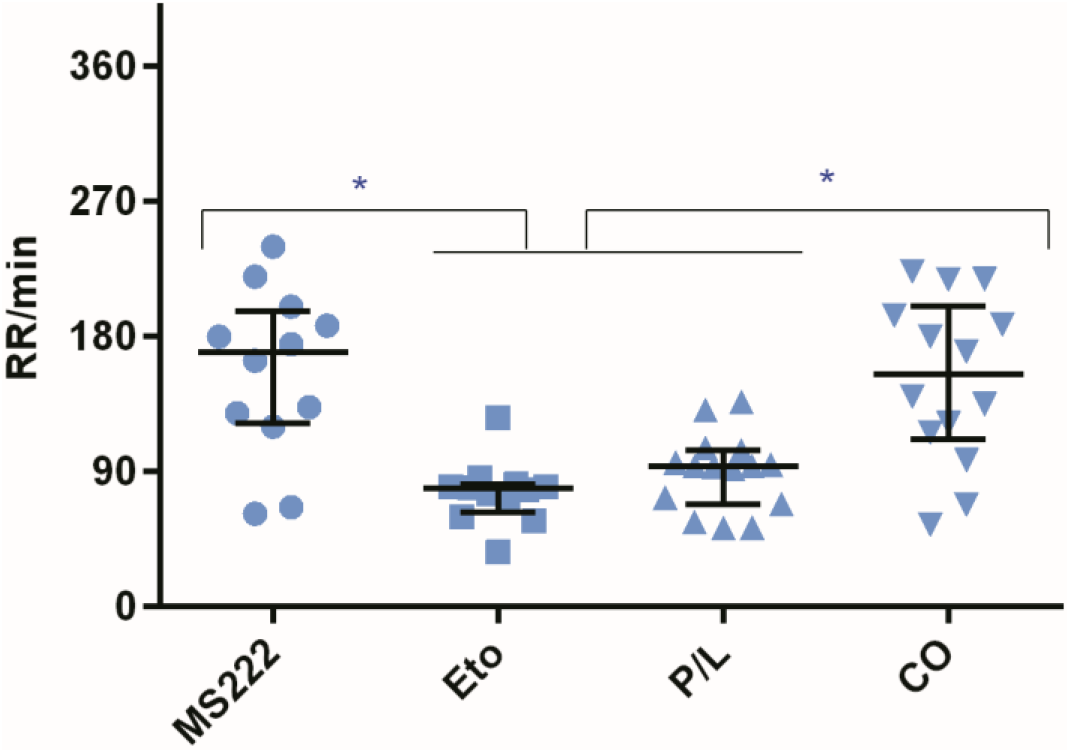
Respiratory rate per minute (RR) of adult zebrafish exposed to different anaesthetics protocols (175mg/L of MS222 (MS222) (n= 14); 2 mg/L of etomidate (Eto) (n= 13); 5mg/L of propofol combined with 150 mg/L of lidocaine (P/L) (n= 15); 45 mg/L of clove oil (CO) (n= 16)); Each point represents an animal. Data are expressed as median [interquartile range]. * p < 0.05

### Novel tank

During the 6 minutes in the novel tank test, no significant differences were found between groups regarding the parameters analysed in the whole apparatus to study long-term recovery of the swimming activity (total distance travelled, average speed, maximum speed) and anxiety-like behaviours potentiated by the exploration of a new environment: angular velocity, number, distance, and duration of erratic movements. This lack of differences between groups was also observed on these parameters analysed in each zone (upper and bottom tank zone). The number of entries in each zone and the indices of time and distance were also similar between groups.

Regarding behavioural comparisons between zones, all groups spent significantly less time in the UP zone compared with the time spent in this zone by chance (180 sec) (*p*≤0.001; Fig. 3A). Also, the time spent (*p*≤0.001; Fig. 3B), distance travelled (*p*≤0.001; Fig. 3C) and maximum speed (*p*<0.05; Fig. 3D) were significantly higher in the bottom of the tank than in the upper zone in all groups. The latency to exit the bottom zone was significantly higher than to exit the upper zone in all groups (Supplementary Table S1).

**FIG. 3.**
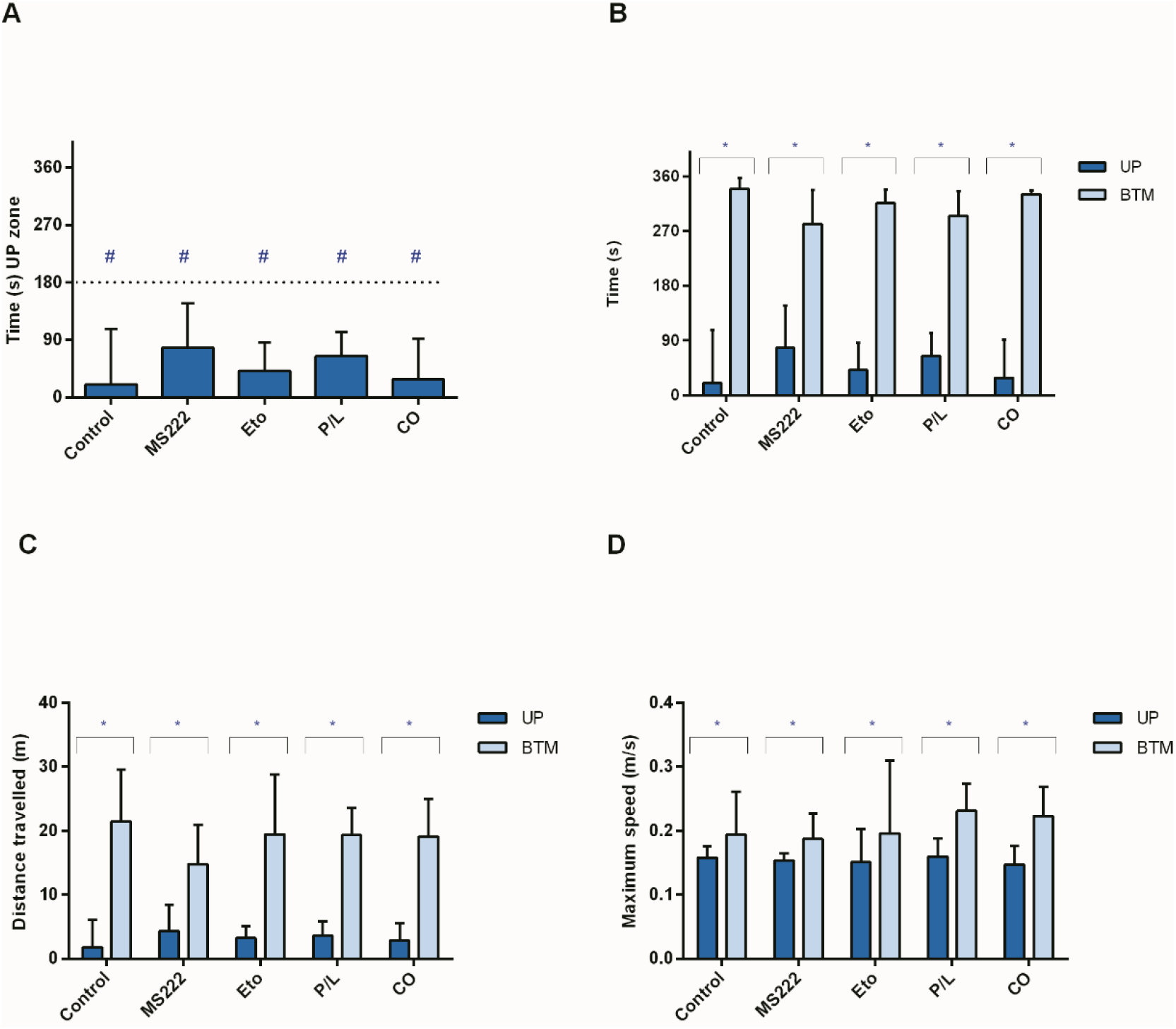
Activity of adult zebrafish in the novel tank for 6 minutes, 28 hours after being exposed to different anaesthetic protocols (175mg/L of MS222 (MS222) (n= 14); 2 mg/L of etomidate (Eto) (n= 15); 5mg/L of propofol combined with 150 mg/L of lidocaine (P/L) (n= 15); 45 mg/L of clove oil (CO) (n= 15)). **(A)** Time spent (s) in the UP zone of the tank; **(B)** Comparison of time spent (s) between zones (UP and BTM); **(C)** Total distance travelled (m) in the UP and BTM zone; **(D)** Maximum speed (m/s) in both zones (UP and BTM). A group of non-anaesthetized animals (n=13) was used as control. UP: upper zone of the tank; BTM: bottom zone of the tank. Data are presented as median [interquartile range]. # p < 0.05 for comparisons between the time spent in the UP zone and the time spent there by chance (180 s); * p < 0.05 for comparisons between zones within treatments.

When the analysis was divided in two segments of 3 minutes, it was observed that all groups spent significantly less time in the UP zone compared with what would be predicted by chance (90 sec) in the first and in the last 3 minutes (*p*≤0.001, Fig. 4A). Also, in both segments of time, all animals spent significantly more time swimming (*p≤*0.001; Fig. 4B or Fig. 4C, respectively) and swam longer distances (*p≤*0.001 or *p*<0.01, respectively, Supplementary Table S1) in the bottom than in the upper zone. Nevertheless, Eto animals increased their swimming time (p=0.011, Fig. 4A), distance and number of entries (p≤0.033, Supplementary Table S1) in the upper zone in the last 3 minutes compared with the first 3 minutes, being the only group to change this behavioural pattern. Also, etomidate-treated animals increased the angular velocity in the whole tank analysis at the last 3 minutes compared with the first 3 minutes (Supplementary Table S1). Other occasional differences in these time segments analysis is reported in the Supplementary Table S1.

**FIG. 4.**
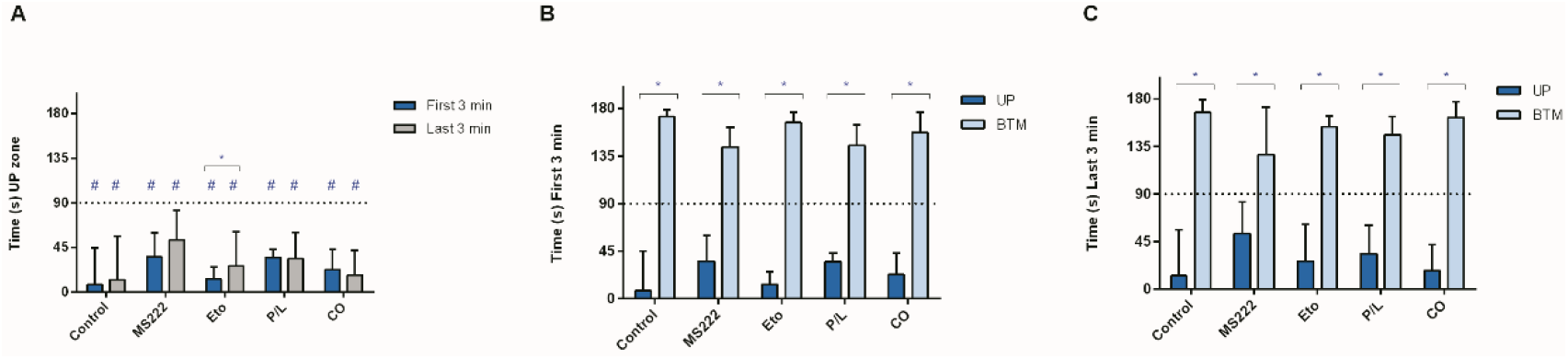
Time spent (s) by adult zebrafish in the novel tank test after exposure to different anaesthetic protocols (175mg/L of MS222 (MS222) (n= 13); 2 mg/L of etomidate (Eto) (n= 15); 5mg/L of propofol combined with 150 mg/L of lidocaine (P/L) (n= 15); 45 mg/L of clove oil (CO) (n= 15)) and results presented by dividing the time in two segments of 3 minutes. **(A)** Comparison of time spent (s) in the UP zone in the first and last 3 minutes; **(B)** Comparison of time spent (s) in each zone (UP and BTM) in the first 3 minutes; **(C)** Comparison of time spent (s) in each zone (UP and BTM) in the last 3 minutes. A group of non-anaesthetized animals (n=13) was used as Control. UP: upper zone of the tank; BTM: bottom zone of the tank. Data are presented as median [interquartile range]. * p < 0.05 for comparisons between zones or between segments of time within treatments; # p < 0.05 for comparisons between the time spent in the UP zone and the time spent there by chance (90 s) in each segment of time.

For both 6-minutes and 3-minutes analysis, none of the groups had an angular velocity significantly higher than 180° s-1, only clove oil-treated animals had a significantly lower value (Supplementary Table S1).

## Discussion

Proper anaesthesia should avoid distress to the animals while inducing immobility and analgesia as needed, and a full recovery in the end. In this study, we tested several anaesthetic protocols for adult zebrafish to determine their efficacy, safety, and recovery quality. In general, independently of anaesthesia protocol, all fish were rendered unconscious, indicated by equilibrium loss, and displayed a full recovery from anaesthesia with no mortality. Also, using the novel tank test, no side-effects on anxiety levels were evident 28 hours after anaesthesia.

MS222 has frequently been referred to as being aversive or causing distress to zebrafish^2, 11, 18, 19^, but our results showed that this anaesthetic was not different from any of the others in terms of erratic movements after anaesthesia. Rapid opercular movements and some erratic movements were also seen during induction in all animals, including the control group, probably for being introduced into a new environment with a small height of water column.

Typically, the optimal anaesthetic should induce a rapid anaesthesia (3 minutes or less) and recovery (5 minutes or less), leave low tissues residues after a withdrawal period of 1 hour or less, have low cost and not be toxic for fish and users^20, 21^. All the anaesthetic protocols tested induced loss of equilibrium in less than 3 minutes. Nevertheless, clove oil induced a quicker loss of the equilibrium compared with propofol/lidocaine. This result may be caused by differences in the anaesthetics solubility and action mechanisms. Propofol is not fully soluble in water and the solution must be vigorously stirred. More importantly, clove oil is highly lipophilic and hence rapidly absorbed through the gills to the bloodstream and transported to the brain and other body tissues^22, 23^, where it acts.

Regarding response to painful stimulus, etomidate did not induce loss of this response. This was expected since etomidate is a hypnotic that does not provide analgesia^9^. Moreover, 45 mg/l of clove oil only caused loss of the response to painful stimuli in one animal (0.67%). Similar findings were observed by Bolasina, de Azevedo ^24^ for guppy, where a 50 mg/l eugenol exposure resulted in light sedation. However, for clove oil, the time under exposure may be the key to achieve analgesia as our preliminary studies showed that clove oil at the same concentration induces analgesia in almost all animals, but it takes longer than 7.5 minutes^25^. Even though 8 P/L animals did not lose the response to painful stimuli (53%), our previous work^13^ showed that propofol/lidocaine at the same dose induced analgesia. In the present study, there was a time limit of 7.5 minutes of anaesthesia after equilibrium loss, thus this result does not mean that P/L cannot produce full anaesthesia in all animals, only that it may takes longer than the period observed. Also, in Valentim, Felix ^13^ study, using egg water instead of system water to prepare the anaesthetics solutions resulted in a higher pH (8.2) than in this study (7± 0.2 pH). The anaesthetic solution pH interferes with its efficacy, as it changes the ratio of ionized and nonionized forms^9^. In the water, the pka of propofol is 11^26^, while the one for lidocaine is 7.7^27^. This increase of the water pH to a value nearer the pka leads to a greater proportion of nonionized forms, resulting in higher bioavailability^28^, and consequent increase in drug efficacy. Thus, the higher pH of the water of the anaesthetic solution in Valentim, Felix ^13^ study, resulted in a quicker anaesthesia and analgesia compared with the present study. Given this, pilot studies to test the effects of water pH in the anaesthetic solutions on animals are recommended, as well as reporting the pH values used, improving data reproducibility and replicability^29^.

MS222 has been described to depress the cardiovascular and respiratory system, interfering with ion regulation^30^ during exposure, especially in long duration procedures. However, it has also been described to stimulate respiratory activity during anaesthesia induction^31, 32^. In the present study, MS222 induced higher opercular movements compared with etomidate and propofol/lidocaine, after the animal was placed in clean water. Thus, the stimulatory mechanism seems to be activated also immediately after the animal has been removed from a short duration anaesthetic bath. CO animals also had higher opercular rates compared with Eto and P/L animals. This was expected as propofol may cause cardiorespiratory depression^33^. In Valentim, Felix ^13^, the opercular rate was higher in the P/L group for the same concentration, probably because the duration of anaesthesia is higher in all animals in the present study. Moreover, some studies^12, 34, 35^ found evidence of ventilation reduction following etomidate exposure. In this situation, animals took several minutes to recover the normal opercular rate^35^, which explains the lower opercular rates in the Eto animals even after they had been placed in clean water for recovery. Regarding clove oil, it has been described to inhibit the respiratory centres in the medulla oblongata^36^, which predicts a decrease in the opercular movements that was not observed in the present study. However, the data from literature is related to the period during anaesthesia and this study shows the opercular rate during the start of anaesthesia recovery.

In general, the temperature at which zebrafish are kept promotes high metabolic rates, thus, once the fish is in clean water, elimination of the anaesthetics is expected to be relatively quick^22^. Indeed, except for one, the animals recovered in less than 10 minutes. The high opercular rate observed in the MS222-treated animals may have promoted a faster regain of the equilibrium compared with the others, as this allowed the anaesthetic to be metabolized and eliminated quickly by the gills. Thus, the induction of MS222 quick recovery is probably due to the local anaesthetic action and stimulatory effects on the respiratory and cardiovascular system^37^. Following this idea, as clove oil-treated animals also had high opercular rates, they may be expected to recover faster. However, the fact that this anaesthetic is an oil and has been described to coat gill epithelia, may make anaesthetic elimination difficult^38^ and thus prolong the anaesthesia.

Our results regarding equilibrium recovery of the P/L group support our previous work^13^. However, Martins, Diniz^12^ reported that this combination induced a more rapid recovery than MS222. This discrepancy in results may be explained by the fact that they used a lower dose of propofol/lidocaine than the one in the present study.

Although MS222 animals took less time to recover the equilibrium, at 1 hour post-anaesthesia, all animals swum in the water column, and anaesthesia groups already exhibited a control level activity. All groups increased distance swum throughout time, showing a normal process of habituation to being placed in a new tank. This is in accordance with previous works^12, 13^, where MS222 and P/L groups recovered at least at 5- and 24-hours post-anaesthesia.

To evaluate anxiety-like behaviours, we used the novel tank test which is based on fish’ natural tendency to react to novelty by initially spending more time at the bottom of the tank and gradually increase their exploration to the upper zone from where threats may appear^39^. We found that fish from all treatments spend more time swim and swam longer distances and at a higher maximum speed in the bottom than in the upper zone of the tank, for all periods of the test, with no differences between groups. Thus, all animals behaved similarly to control levels, showing no alterations on the anxiety profile. Apart from minimal scattered differences, etomidate group in the last 3 minutes of analysis showed an increase of time, distance, and number of entries in the UP zone compared with the first 3 minutes. Although these increases are not enough to characterize an anxiolytic-like behaviour, this data may be related to the properties of etomidate to block cortisol production^40^, reducing stress responses and facing the upper zone of the novel tank with less cautious throughout time quicker than the other groups. Etomidate-treated animals also exhibited an increase in angular velocity from the first to the last 3 minutes. An increase in angular velocity is often associated with escape turns indicative of stress. However, the angular velocity per group was never above 180° s^-1^, which indicates routine turns^17^.

The present study shows that all protocols are effective for anaesthetising adult zebrafish, without mortalities or significant alterations on behaviour and with a quick recovery of normal activity (at least within an hour of anaesthesia administration). Although the behavioural alterations induced by etomidate in the novel tank were minimal and do not indicate a different anxiety profile, its use must be considered carefully depending on the experimental objective, as its effect on cortisol release may be an unwanted interference with research results. The choice of the best anaesthetic protocol will also depend on the procedure invasiveness and duration. If a procedure requires analgesia, MS222 and propofol/lidocaine protocol will be the most adequate, although P/L animals may take longer to achieve analgesia with these concentrations. Further studies should be done with animals from other genetic backgrounds and life stages to gather more information regarding the potential impact of different anaesthetic protocols in the zebrafish and to refine the concentrations needed for each procedure.

## Acknowledgements

The authors would like to thank to the i3s Zebrafish Facility staff for helping with the maintenance of the tanks and the animals in ideal conditions.

## Authorship confirmation statement

AMV conceived the experimental design. JF and AMV conducted the experiments. SJ analysed the data and wrote the first version of the manuscript. AMV gave important input and close supervision to the manuscript writing. AO provided critical feedback. All authors discussed the results, contributed to the final manuscript, and approved the manuscript for publication.

## Supplementary Material

Supplementary Figure S1

Supplementary Table S1

