## Supplemental data for "Adult zebrafish anaesthesia: a study of efficacy and behavioural recovery of different anaesthetics"

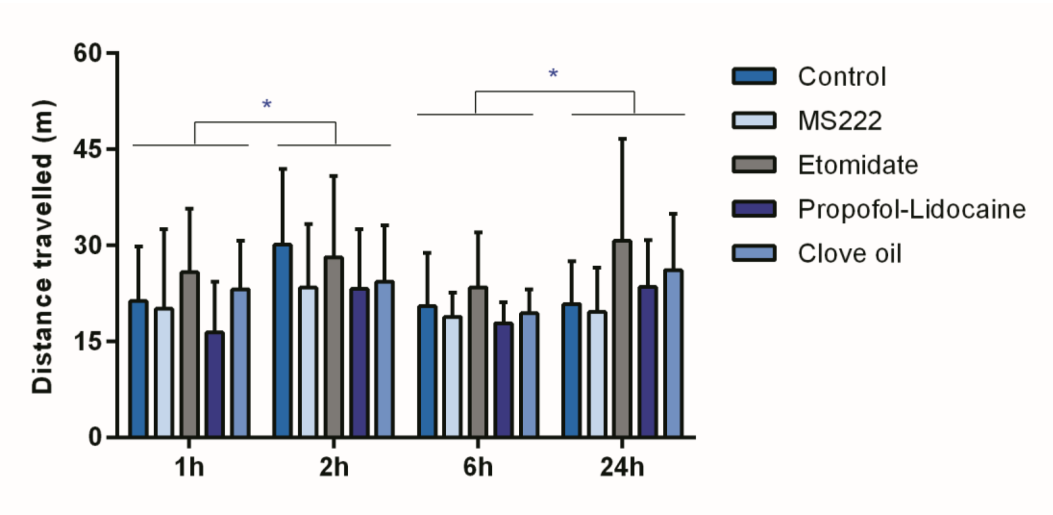


**FIG. S1.** Distance travelled by adult zebrafish 1, 2, 6 and 24h after treatment with different anaesthetics protocols (175mg/L of MS222 (MS222) (n= 13); 2 mg/L of etomidate (Eto) (n= 11); 5mg/L of propofol combined with 150 mg/L of lidocaine (P/L) (n= 12); 45 mg/L of clove oil (CO) (n= 15). A group of non-anaesthetized animals (n=12) was used as Control. Data are expressed as median [interquartile range]. * $p < 0.05$

Table S1. Significant results from the novel tank test that are not represented in the graphs.

| Variable | Description | Treatment | Test for comparison between samples | P-value | Statistical test value[a] | Median [IQR] or Mean ± SD |
|---|---|---|---|---|---|---|
| Angular velocity (° s-1) | First 3min vs Last 3min | Eto | Paired t-test | 0.003 | t(14)= -3.53 | First 3min: 169.1 ± 40.2<br>Last 3min: 196.8 ± 47.8 |
| Angular velocity (° s-1) | First 3min | CO | One-Sample Test | 0.040 | t(14)= - 2.267 | First 3min: 161.1 ± 32.34 |
| Distance | UP (First 3min vs Last 3min) | Eto | Wilcoxon signed rank | 0.023 | 2.272 | First 3min: 0.97 [1.49]<br>Last 3min: 2.76 [2.41] |
| Number of entries | UP (First 3min vs Last 3min) | Eto | Wilcoxon signed rank | 0.033 | 2.137 | First 3min: 8 [8]<br>Last 3min: 16 [17] |
| Average Speed (m/s) | UP (First 3min vs Last 3min) | Control | Paired t-test | 0.028 | t(10)= - 2.574 | First 3min: 0.06 ± 0.03<br>Last 3min: 0.09 ± 0.04 |
| Maximum speed (m/s) | UP (First 3min vs Last 3min) | Control | Paired t-test | 0.009 | t(10)= - 3.247 | First 3min: 0.10 ± 0.06<br>Last 3min: 0.15 ± 0.04 |
| Distance Travelled (m) | BTM (First 3min vs Last 3min) | MS222 | Wilcoxon signed rank | 0.006 | -2.760 | First 3min: 8.5 [5.05]<br>Last 3min: 7.09 [5.05] |
| Time (s) | BTM (First 3min vs Last 3min) | Eto | Wilcoxon signed rank | 0.011 | -2.556 | First 3min: 166.7 [21.60]<br>Last 3min: 153.6 [44.80] |
| Number of entries | BTM (First 3min vs Last 3min) | Eto | Wilcoxon signed rank | 0.044 | 2.017 | First 3min: 8 [7]<br>Last 3min: 16 [18] |

| | | | | | | |
|---|---|---|---|---|---|---|
| Latency to 1st exit (s) | BTM (First 3min vs Last 3min) | Control | Wilcoxon signed rank | 0.028 | -2.201 | First 3min: 23.75 [37.30] Last 3min: 9 [134.90] |
| Latency to 1st exit (s) | BTM (First 3min vs Last 3min) | P/L | Wilcoxon signed rank | 0.028 | -2.197 | First 3min: 58.05 [69.07] Last 3min: 9.7 [18.95] |
| Distance (m) | UP vs BTM First 3min | Control | Wilcoxon signed rank | 0.001 | 3.180 | UP First 3min: 0.75 [0.29] BTM First 3min: 11.12 [8.39] |
| Distance (m) | UP vs BTM First 3min | MS222 | Paired t-test | 0.001 | t(12)=- 4.295 | UP First 3min: 2.58 ± 2.24 BTM First 3min: 9.52 ± 4.74 |
| Distance (m) | UP vs BTM First 3min | Eto | Wilcoxon signed rank | 0.001 | 3.408 | UP First 3min: 0.97 [1.49] BTM First 3min: 9.63 [6.31] |
| Distance (m) | UP vs BTM First 3min | P/L | Paired t-test | 0.000 | t(14)= - 8.536 | UP First 3min: 1.70 ± 0.97 BTM First 3min: 10.02 ± 3.11 |
| Distance (m) | UP vs BTM First 3min | CO | Paired t-test | 0.000 | t(14)= - 8.118 | UP First 3min: 1.71 ± 1.45 BTM First 3min: 9.92 ± 3.09 |
| Latency to 1st exit (s) | UP vs BTM First 3min | Control | Wilcoxon signed rank | 0.012 | 2.521 | UP First 3min: 0.6 [0.75] BTM First 3min: 23.75 [37.30] |
| Latency to 1st exit (s) | UP vs BTM First 3min | MS222 | Paired t-test | 0.000 | t(10)= - 5.414 | UP First 3min: 1.01 ± 0.35 BTM First 3min: 28.37 ± 16.75 |
| Latency to 1st exit (s) | UP vs BTM First 3min | Eto | Wilcoxon signed rank | 0.001 | 3.296 | UP First 3min: 0.7 [0.40] BTM First 3min: 29.6 [80.33] |

| | | | | | | |
|---|---|---|---|---|---|---|
| Latency to 1st exit (s) | UP vs BTM First 3min | P/L | Wilcoxon signed rank | 0.001 | 3.296 | UP First 3min: 0.7 [0.70]<br>BTM First 3min: 58.05 [69.07] |
| Latency to 1st exit (s) | UP vs BTM First 3min | CO | Wilcoxon signed rank | 0.002 | 3.059 | UP First 3min: 1.0 [0.40]<br>BTM First 3min: 34.8 [47.05] |
| Distance (m) | UP vs BTM Last 3min | Control | Paired t-test | 0.001 | t(12)= - 6.019 | UP Last 3min: 1.74 ± 1.90<br>BTM Last 3min: 10.06 ± 4.04 |
| Distance (m) | UP vs BTM Last 3min | MS222 | Wilcoxon signed rank | 0.007 | 2.691 | UP Last 3min: 2.69 [4.53]<br>BTM Last 3min: 7.09 [5.05] |
| Distance (m) | UP vs BTM Last 3min | Eto | Paired t-test | 0.000 | t(14)= - 5.525 | UP Last 3min: 2.31 ± 1.32<br>BTM Last 3min: 9.73 ± 4.22 |
| Distance (m) | UP vs BTM Last 3min | P/L | Paired t-test | 0.000 | t(14)= - 5.612 | UP Last 3min: 1.96 ± 1.38<br>BTM Last 3min: 9.32 ± 4.25 |
| Distance (m) | UP vs BTM Last 3min | CO | Wilcoxon signed rank | 0.001 | 3.237 | UP Last 3min: 1.46 [2.25]<br>BTM Last 3min: 9.20 [6.85] |
| Average speed (m/s) | UP vs BTM Last 3min | MS222 | Paired t-test | 0.021 | t(12)= 2.657 | UP Last 3min: 0.07 ± 0.02<br>BTM Last 3min: 0.06 ± 0.02 |
| Average speed (m/s) | UP vs BTM Last 3min | Eto | Wilcoxon signed rank | 0.048 | - 1.979 | UP Last 3min: 0.07 [0.06]<br>BTM Last 3min: 0.06 [0.04] |
| Average speed (m/s) | UP vs BTM Last 3min | CO | Wilcoxon signed rank | 0.044 | - 2.014 | UP Last 3min: 0.06 [0.04]<br>BTM Last 3min: 0.06 [0.03] |

CO: Clove Oil; Eto: Etomidate; P/L: Propofol- Lidocaine; UP: the upper zone of the novel tank; BTM: the bottom zone of the novel tank.

[a] For paired Students' t-test: t(df)= t value. For Wilcoxon signed rank is presented the standardized statistical test value.
